# The left angular gyrus is causally involved in information buffering and context formation: evidence from a narrative reading task

**DOI:** 10.1101/841478

**Authors:** Francesca M. Branzi, Gorana Pobric, JeYoung Jung, Matthew A. Lambon Ralph

**Author notes:** **Address for correspondence:** Dr. Francesca M. Branzi, E. or Prof. Matthew A. Lambon Ralph, E.

## Abstract

The role of the left angular gyrus (Ag) in semantic processing remains unclear. In this study, we used transcranial magnetic stimulation (TMS) to test the hypothesis that the left Ag supports the information buffering processes necessary for context-dependent semantic integration (narrative reading task). We applied online TMS to the left Ag to disrupt the buffering processes while female and male human participants integrated information between two paragraphs of text presented sequentially (i.e., context and target conditions). We assessed the effect of TMS on the left Ag by recording reading times for target conditions during the reading task and by asking participants to retrieve contextual information given the target condition as cue in a successive memory task. TMS applied over the left Ag during the reading task impaired the retrieval of contextual information in the memory task without affecting reading times. These results suggest that the Ag supports information buffering and context formation. This study provides the first evidence of a causal role of the left Ag in language tasks that require context-dependent semantic integration.

**Significance statement:** The present study offers novel insights into the processes that are associated with the left angular gyrus (Ag) activation during context-dependent naturalistic language processing. We used TMS while human participants read written narratives to interfere with online language integration. We measured how TMS over the left Ag was affecting reading times and retrieval of integrated narrative representations. We were able to show that the left Ag is causally involved in information buffering and context formation.

## 1. Introduction

The left angular gyrus (Ag) is implicated in a range of cognitive activities (Seghier, 2013; Humphreys and Lambon Ralph, 2015). Amongst these, its specific role during semantic tasks is still under debate. Some have proposed that the left Ag might play a key role in the integration of semantic information (Humphries et al., 2007; Binder et al., 2009), while others have argued that left Ag supports buffering processes that operate on content that is integrated elsewhere (Vilberg and Rugg, 2012; Humphreys and Lambon Ralph, 2015, 2017). The latter hypothesis is in accord with evidence showing that left Ag is engaged when semantic tasks require context-dependent integration (Simony et al., 2016; Baldassano et al., 2017; van der Linden et al., 2017; Branzi et al., 2019), but deactivated when tasks do not require such process (Humphreys et al., 2015; Humphreys and Lambon Ralph, 2015). This transcranial magnetic stimulation (TMS) study tested the hypothesis that, during semantic tasks, the primary functional role of the left Ag reflects information buffering and context formation.

TMS was applied over the left Ag while human participants read narratives composed of two consecutive passages (context and target conditions). We hypothesised that if the left Ag is necessary for integrating context and target information, Ag TMS applied between context and target presentation should affect the encoding of a context-target integrated representation. To test this, after the reading task, participants performed a memory task. Importantly, Ag activity is positively engaged during encoding and predictive of better performance at memory retention only when the buffered information fits with a knowledge-based schema (Humphreys and Lambon Ralph, 2015; van der Linden et al., 2017). Hence, an Ag TMS effect should be observed only when context and target passages are semantically coherent (i.e., same schema), but not when their integration requires context updating. To test this, Ag TMS was applied during two different types of narrative conditions in which the same target paragraph was preceded by different types of context: (i) a highly congruent context (HC) which maximised the information contained in a coherent semantic gestalt; (ii) a low–congruent (LC) paragraph with a divergent meaning, thus requiring updating the semantic context. In the memory task, participants were presented with the target condition (both HC and LC) as a cue and required to retrieve context-related information (“Cue Target” trials). We expected Ag TMS to impair behavioural performance in HC conditions, but to have no effect on the retrieval of LC conditions.

Furthermore, memory representations that include elements following and preceding the update of information, are harder to retrieve than representations that require retrieval of associations within the same event (Speer and Zacks, 2005; Swallow et al., 2009). Accordingly, we expected to find increased response times (RTs) and error rates for low over high context-to-target congruency under standard conditions (i.e., no Ag TMS) and that Ag TMS would diminish this effect given the Ag’s role in information buffering and context formation. To rule out the possibility that TMS was affecting encoding in general, rather than the encoding of context-target integrated representations specifically, we employed a control condition in the memory task. In “Cue Context trials”, participants were required to retrieve the same context-related information as in the Cue Target conditions. However, they were provided with some contextual information as cue.

Finally, if the left Ag is necessary for integrating information between context and target, Ag TMS between context and target presentation during reading should affect also reading times locked to the target presentation. Similarly as in the memory task, we expected this effect for HC conditions only (van der Linden et al., 2017). Besides HC and LC conditions, in the reading task we also included (iii) a no context control condition (NC) in which the same target was preceded by a number reading task. This control condition allowed verification that TMS effects measured during the reading task were attributable to the buffering of contextual information only.

## 2. Materials and Methods

### 2.1. Participants

Eighteen volunteers took part in the study (average age = 22, standard deviation (SD) = 3; N female = 12). All participants were right-handed (Oldfield, 1971), native English speakers with no history of neurological or psychiatric disorders and normal or corrected–to–normal vision. Written informed consent was obtained from all participants. The experiment was approved by the local ethics committee.

### 2.2. Stimuli

#### Reading task

The experimental stimuli used in the reading task were the same as in Branzi et al. (2019). Thus, a total of 40 narrative pairs, each one composed by two paragraphs, were employed in the reading task. For each narrative pair, the same second paragraph (target) was preceded by different first paragraphs (contexts) that could be either high–congruent (i.e., HC) or low–congruent (i.e., LC) with the target in terms of meaning. Both HC and LC context paragraphs could be integrated with the targets paragraphs, though a reworking of the evolving semantic context was required after LC contexts, because of a shift in the semantic context (see **Table 1** for an example of the stimuli; for the full list of the stimuli used in the reading task see Branzi et al., 2019). Homonym words (e.g., bank), presented at the beginning of the target paragraph, were employed to determine the exact point in the paragraph in which the shift in the semantic context should have been experienced. Finally, the NC condition, where the target (the same as in HC and LC conditions) was preceded by a string of numbers, was employed as a control condition. In fact, the NC condition allowed to verify that any effect of TMS measured for HC and LC conditions was due to buffering processes, and not to bottom-up attention triggered by the presentation of the target paragraph.

**Table 1.**
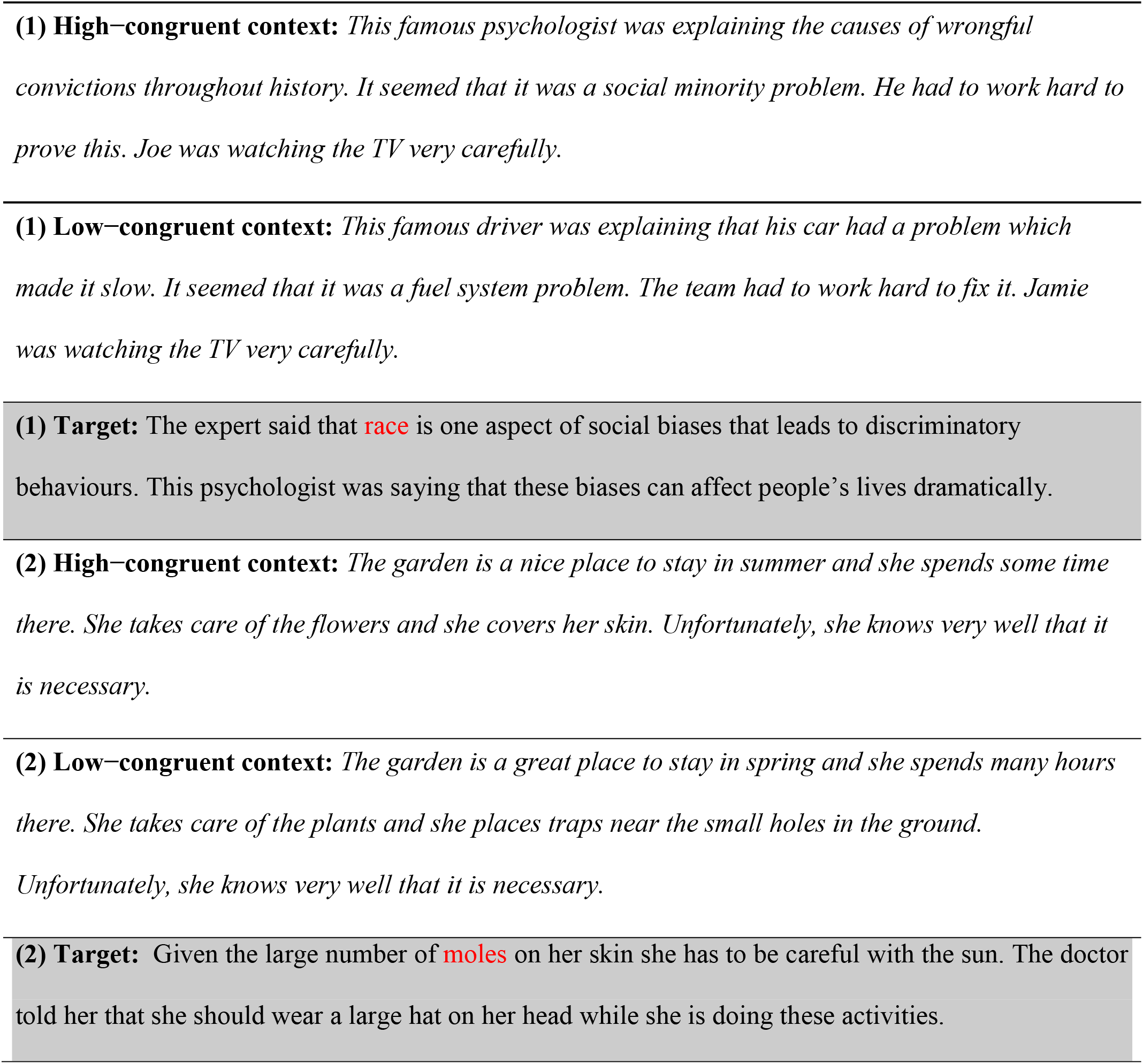
Examples of stimuli for HC and LC conditions (Context and Target paragraphs). The shift of semantic context after LC contexts was expected to be perceived after the critical ambiguous word, here depicted in red.

#### Memory task

The experimental stimuli employed in the memory task were 75 items, consisting in (1) the presentation of a “cue” displayed on the screen followed by (2) the presentation of a question relative to the context part of the narrative along with three possible answers (Q&As, see **Table 2** for some examples). Importantly, in the memory task there were different types of conditions. These conditions did not differ in respect to the type of information that participants had to retrieve (the same contextual information), rather they differed in the type of cue. In one condition the cues only corresponded to the information in the target paragraph [i.e., HC Cue Target conditions (n = 25) and LC Cue Target conditions (n = 25)], and hence tested participants’ ability to retrieve target-context integrated representations. By contrast in the other condition, the cues contained contextual information [i.e., Cue Context conditions (n = 25)] and thus did not require participants to retrieve the integrated meaning of the two sequential paragraphs. The type of contextual information that participants were required to retrieve during the memory task corresponded to a variety of episodic details (e.g., who, where, when) (see **Table 2**).

**Table 2.**
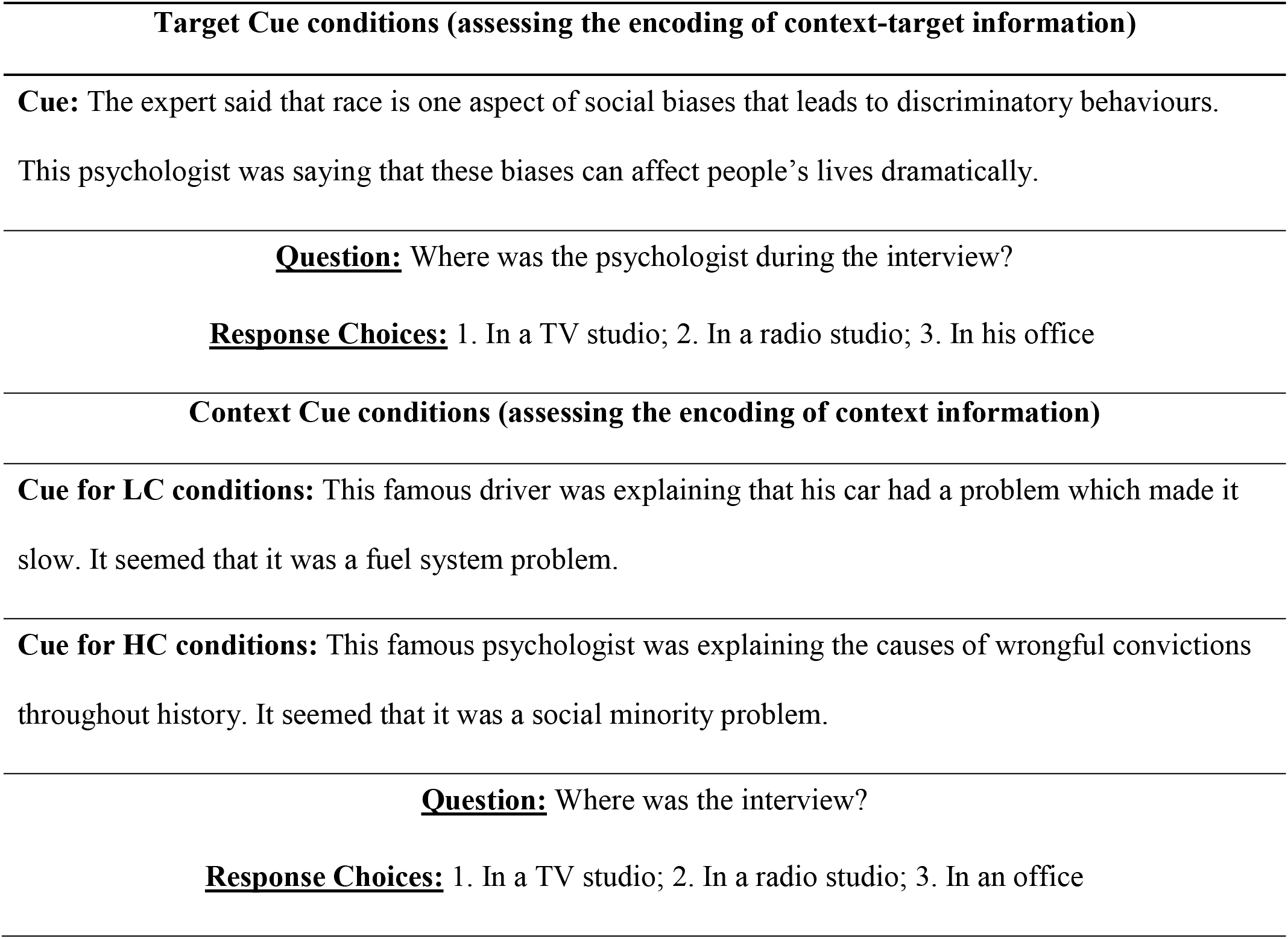
Example of stimuli for Target Cue and Context Cue conditions, assessing the encoding of context-target and context information, respectively. The stimuli presented below refer to the narrative number 1 presented in Table 1.

### 2.3. Task procedures

Before starting the experimental study, all participants signed an informed consent and were given written and oral instructions. A PC running ePrime (Psychology Software Tools, Inc., Pittsburgh, PA) was used to present the items and record participants’ behavioural responses. The participants completed three sessions on different days. During two out of three sessions, participants received TMS (TMS over the left Ag and TMS over the Vertex). In one session, TMS was not used. The sessions were at least 1 week apart (average days between sessions = 12, SD = 3) and the order of sessions was counterbalanced across participants.

In each session, each participant performed two different tasks: a reading task and a memory task, both divided in two parts. Specifically, each session started with a reading task, in which participants had to read half of the narratives. The reading task was immediately followed by a memory task, comprised of questions about the narratives presented in the reading task. After a short break (10 minutes), the remaining narratives were presented in a second reading task, which was followed by another memory task (including questions relating to the narratives presented in the second reading task).

During the TMS sessions, online TMS was delivered during each trial in the reading tasks (for detail see **Figure 1** and *Stimulation Parameters and Stimulation Sites*). TMS was not applied during the memory tasks. We did not apply TMS during the memory task since the goal of the present study was to measure the contribution of the left Ag for semantic integration during comprehension. Since the left Ag is implicated not only in language processing but also in memory retrieval (Rugg and Vilberg, 2013; Humphreys and Lambon Ralph, 2015), not applying TMS during the memory task ensured that any observed TMS effect reflected the contribution of Ag for integrating semantic information during the reading task.

**Figure 1.**
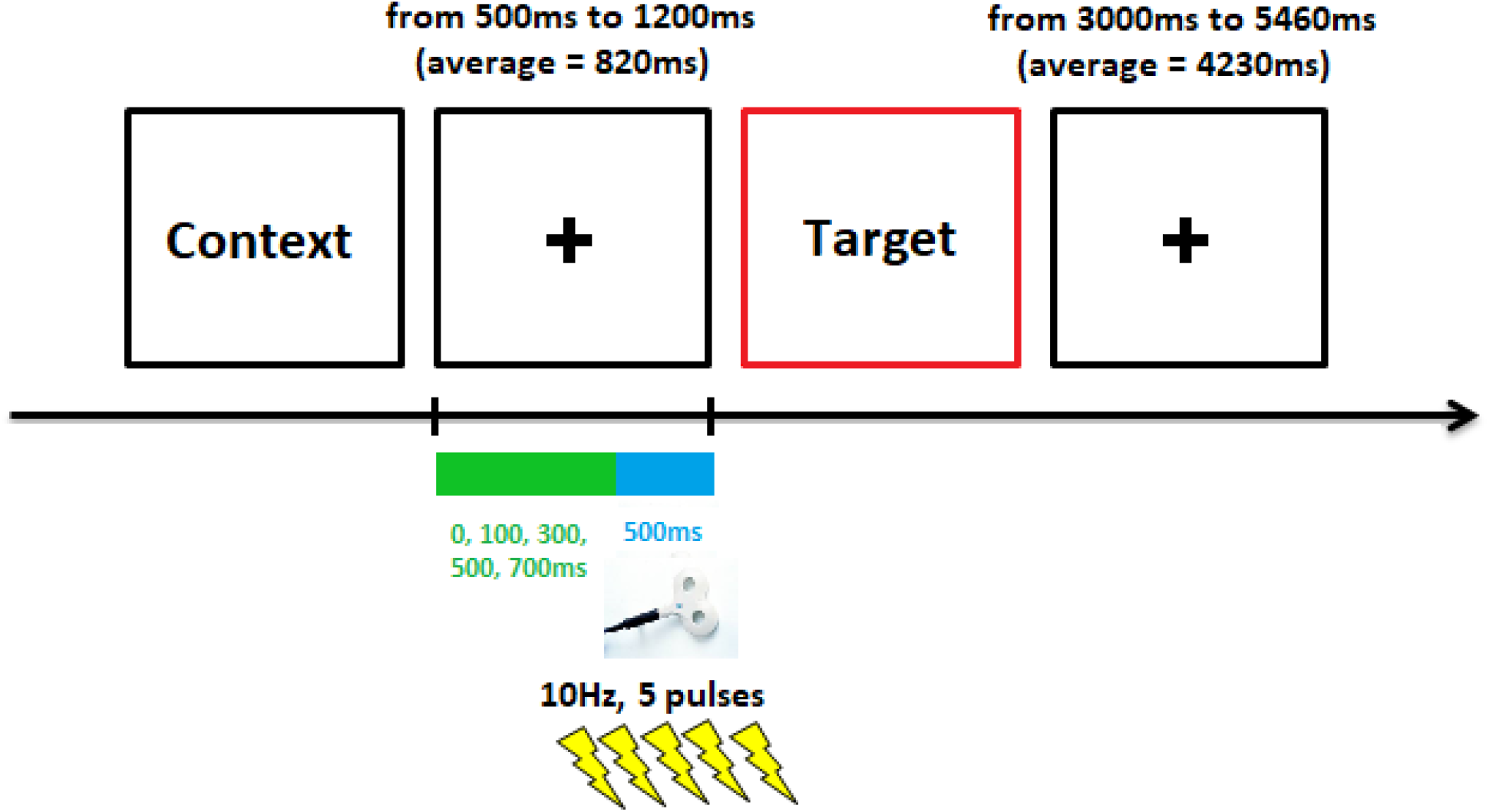
Time line of trial presentation during the reading task.

At the end of each TMS session, participants completed a questionnaire in which they reported the extent to which TMS was perceived as uncomfortable and distracting (scales from 1 = not very, to 7 = very). The scores obtained were overall low (Ag distracting: average = 3.8, SD = 1.3; Ag uncomfortable: average = 3.9, SD = 1.3; Vertex distracting: average = 3.3, SD = 1.2; Vertex uncomfortable: average = 3, SD = 1.4), suggesting that the TMS protocol induced only minimum discomfort and distractibility during the task. A trend toward significance indicated that overall TMS over the Ag obtained slightly higher scores on both scales [main effect of Site: F (1, 17) = 4.185, p = 0.057, *η*p^2^ = 0.198]. Finally, lack of significant Site × Type of Scale interaction [F (1, 17) = 1.034, p = 0.323, *η*p^2^ = 0.057] suggested that TMS on both sites was inducing similar ratings for discomfort and distractibility.

#### Memory task

There were 25 items per condition (Cue Context, LC Cue Target and HC Cue Target). The LC Cue Target trials (25) were presented with half of the Cue Context trials (12 or 13), always after reading the LC narratives. The HC Cue Target trials (25) were presented with the remaining Cue Context trials (13 or 12), always after the HC narratives. The memory task was a 3 alternative-forced-choice task. Each trial started with the presentation of a “cue” displayed on the screen until participants made a button response, which was followed by the presentation of a question and 3 possible responses (Q&Rs). The Q&Rs were displayed until the participants made their selection by button response, up to a time limit of 8.5 seconds (s). Rest time (black fixation cross) was presented between cue and Q&Rs and between trials (fixed time intervals: 250ms and 2000ms, respectively). Finally, whilst the stimuli presented in the reading task were the very same across all the three sessions, those presented during the memory task were always different across the three sessions.

#### Reading task

In the reading task, there were 40 items per condition (HC, LC and NC). As in our previous study (Branzi et al., 2019), each trial consisted in the presentation of two paragraphs of text (context and target) that participants had to read silently (verbal material and numbers). Contexts and targets were displayed on the screen until participants pressed a button to indicate that they had finished reading the paragraph (for both contexts and targets). The instruction emphasized speed, given the limited amount of time for reading [max. duration for context was 12s and for targets was 7.5s], but also the need to understand and encode the information presented in the narratives. We informed participants that at the end of each reading task they would perform a memory task, requiring them to answer questions on the content of the narratives. We also specified that in order to perform the task it would be necessary to integrate the information presented in contexts and targets. Rest time was varied between context and targets [range between 500 milliseconds (ms) and 1200ms, average time = 820ms] and between trials (range between 3000ms and 5460ms, average time = 4230ms) during which a black fixation cross was presented.

HC and LC trials were presented separately. Given that HC and LC conditions were terminating with the same target paragraph, mixing them during the reading task would have created confusion during the memory task. Thus, the first reading task in each session included either HC trials (40) or LC trials (40), and half of the NC trials (20). The second reading task included the remaining narratives (either LC trials or HC trials, and the remaining NC trials). The order of presentation of LC and HC conditions was counterbalanced across participants and sessions.

#### Stimulation Parameters and Stimulation Sites

TMS was delivered using a Magstim Rapid2 stimulator (Magstim Co., Whitland, UK) and a figure-of-eight coil with a diameter of 70 mm. Stimulation was performed at 120% of the individual's motor threshold, measured before the start of the first session (mean stimulation intensity = 71, SD = 8, range = 54 - 84). The resting motor threshold of the relaxed contralateral abductor pollicis brevis muscle was measured as the lowest stimulation intensity able to cause a visible twitch in the muscle 5 out of 10 times (Sandrini et al., 2011).

For each trial, one train of five pulses (10Hz for 500ms) was delivered. This stimulation protocol has already been used over the parietal cortex to induce inhibitory effects during language processing (Sliwinska et al., 2015). Unlike previous studies, the first pulse was administered before the presentation of the target (see **Figure 1**), to avoid disrupting reading (TMS to the Ag can induce eye twitches). A pilot study confirmed that this procedure was successful for obtaining the expected TMS effects. The rest time between the context and target phases was variable and randomized (**see Figure 1**). Thus, participants were not able to anticipate when they would receive TMS. Finally, the TMS frequency, intensity and duration were well within established international safety limits (Rossini et al., 2015).

The stimulation site for the left Ag region corresponded to the MNI coordinates (x = −48, y = −63, z = 36) derived from our previous fMRI study, in which the same experimental material was used and where Ag activity was modulated by context integration (Branzi et al., 2019) (see **Figure 2**). During the Ag TMS testing session, a Polaris Vicra infrared camera (Northern Digital, Waterloo, ON, Canada) was used in conjunction with the Brainsight frameless stereotaxy system (Rogue Research, Montreal, QC, Canada) to register the participant's head to their own MRI scan to accurately target stimulation throughout the experiment. The location of the control site, i.e., Vertex, was established for each participant by using the international 10-20 system (Steinmetz et al., 1989). The halfway intersection of the two lines was marked using a skin marker and corresponded to individual Vertex control site.

**Figure 2.**
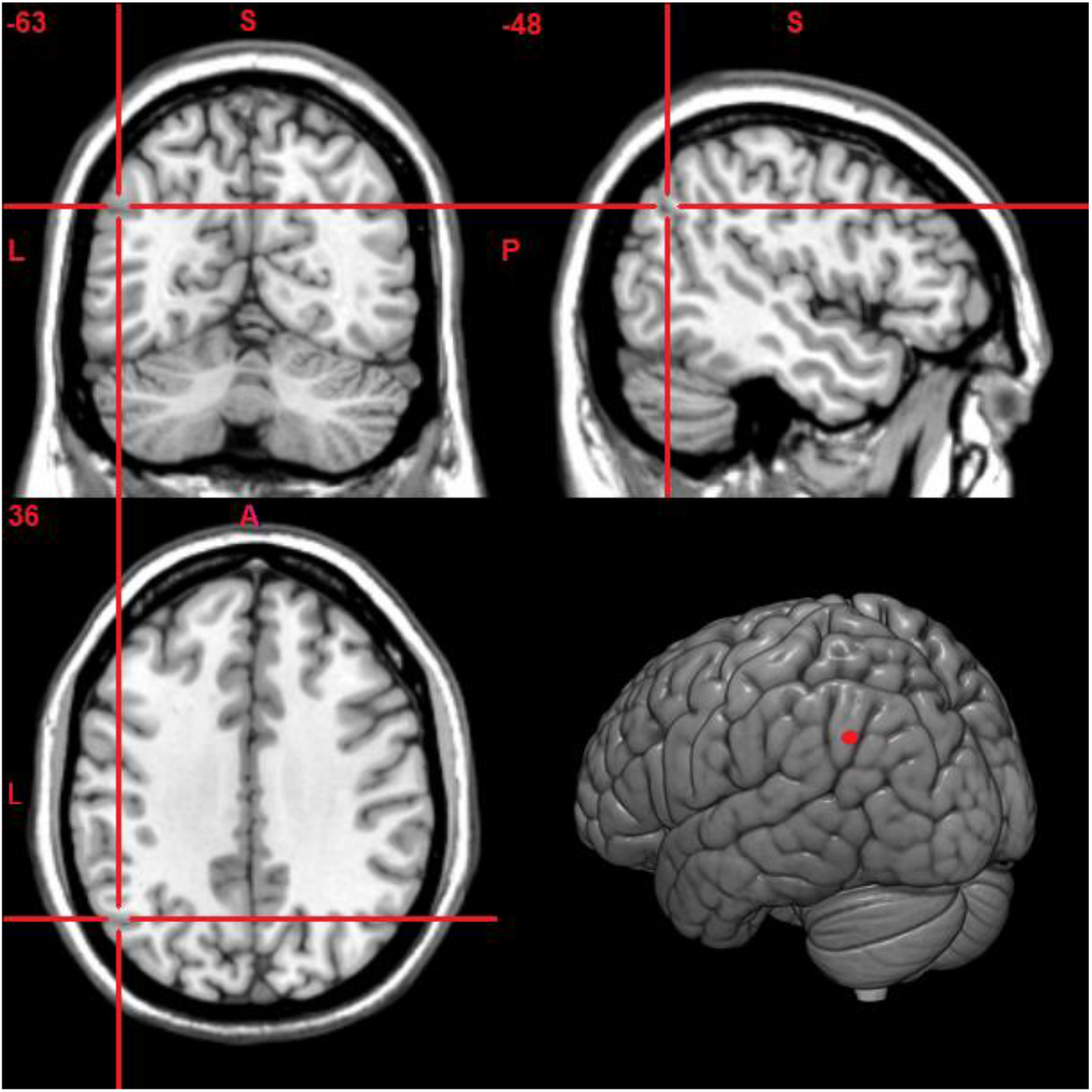
The left Ag stimulation site pinpointed on the MNI cortical template.

### 2.4. Data analysis

#### Behavioural data analyses

##### Memory Task

Behavioural analyses were performed on RTs and accuracy measures. RT analysis was conducted only for correct trials. Moreover, after having eliminated error trials, also trials exceeding 3 SDs above or below a given participant's mean were excluded from the RT analyses, causing an overall loss of 2% of data, across all task conditions and type of sessions. Then, we conducted two separate 3 × 3 within-subject ANOVAs (one for RTs and one for accuracy measures) with the repeated-measures factors condition (NC, LC and HC) and session-type (no TMS, TMS to left Ag, and TMS to vertex). Bonferroni correction for multiple comparisons was applied on post hoc pairwise contrasts.

##### Reading Task

Behavioural analyses were performed on reading times only. In fact, since participants were reading silently, in this task it was not possible to measure error rates. First, trials exceeding 3 SDs above or below a given participant's mean were excluded from the analyses, causing a loss of 0.2% of data, across all task conditions and type of sessions. Then, the effect of TMS on the left Ag was assessed conducting a 3 × 3 within-subject ANOVA with the repeated-measures factors condition (NC, LC and HC) and session-type (no TMS, TMS to left Ag, and TMS to vertex). Bonferroni correction for multiple comparisons was applied on post hoc pairwise contrasts.

## 3. Results

### 3.1. Memory task

RTs showed a main effect of condition [F (2, 34) = 9.443, p = 0.001, *η*p^2^ = 0.357]. Post-hoc comparisons indicated that speed for LC Cue Target condition was slower as compared HC Cue Target (p = 0.003) and Cue Context conditions (p = 0.009). There was no significant difference in speed between the HC Cue Target and Cue Context conditions (p > 0.999). The main effect of session type was not significant [F (2, 34) = 0.188, p = 0.830, *η*p^2^ = 0.011], suggesting that overall performance was not particularly affected by TMS. However, the results revealed a significant condition × session-type interaction [F (4, 68) = 3.446, p = 0.013, *η*p^2^ = 0.169] that we explored via planned pairwise comparisons. We hypothesised that, in absence of TMS on Ag, the need to update semantic context in LC Cue Target conditions would have induced slower RTs as compared to HC Cue Target conditions. In keeping with this prediction, we found that LC Cue Target conditions were significantly slower than HC Cue Target conditions when TMS was applied to the vertex (control site) [t (17) = 3.779, p = 0.003 (adjusted p-value), *d*′= 0.89] or when it was not applied at all [t (17) = 2.613, p = 0.05 (adjusted p-value), *d*′= 0.61] (see **Figure 3**). We also hypothesised that Ag TMS would eliminate this effect by affecting the HC Cue Target conditions specifically. In accord with our predictions, the LC>HC effect was not observed when TMS was applied to the left Ag [t (17) = 0.227, p > 0.999 (adjusted p-value), *d*′= 0.053]. This was likely due to fact that HC Cue Target conditions became slower when TMS was applied on the left Ag than when it was delivered to the vertex [t (17) = 2.463, p = 0.05 (adjusted p-value), *d*′ = 0.58] or when it was not delivered at all [t (17) = 2.576, p = 0.04 (adjusted p-value), *d*′= 0.607]. Finally, TMS to the left Ag did not impair RT performance for LC Cue Target conditions [all ps > 0.999] or context cue conditions [all ps > 0.42].

**Figure 3.**
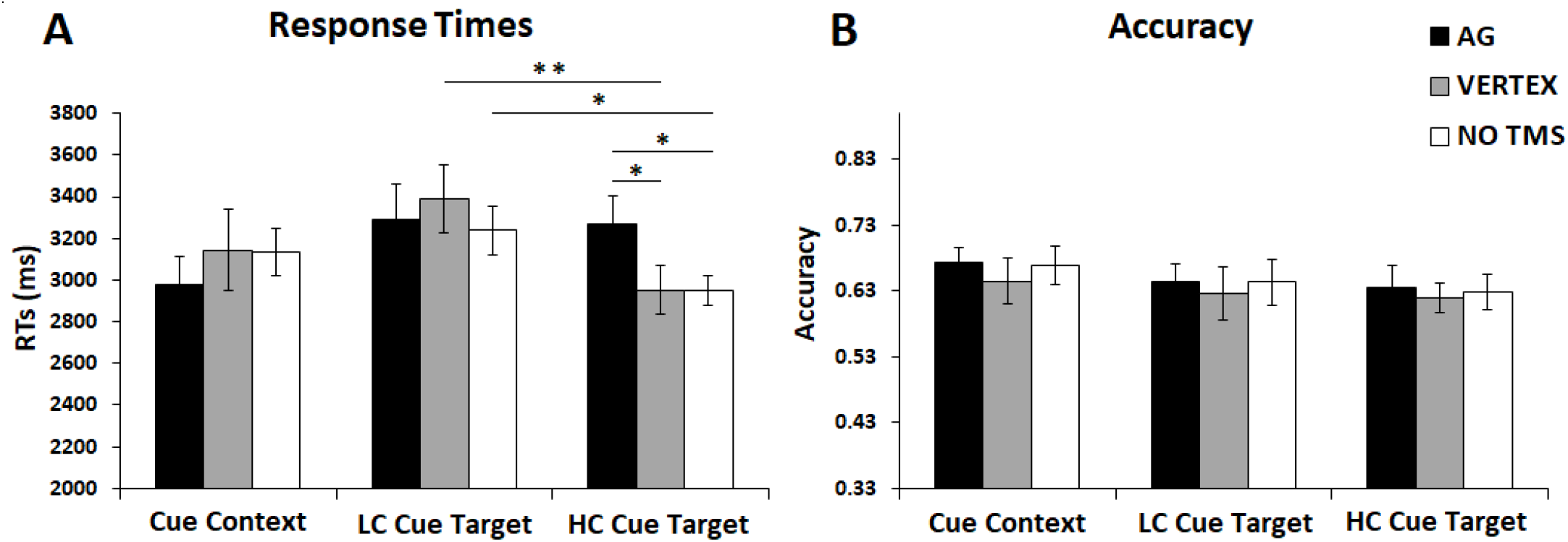
Memory task. Results for (A) RTs and (B) Accuracy (proportion of correct responses) for Cue Context, LC Cue Target and HC Cue Target conditions. Error bars correspond to Standard Errors (SEs).

The accuracy data did not reveal any significant effects (main effect of condition [F (2, 34) = 2.087, p = 0.140, *η*p^2^ = 0.109]; main effect of session-type [F (2, 34) = 0.515, p = 0.602, *η*p^2^ = 0.029]; condition × session-type interaction [F (4, 68) = 0.030, p = 0.998, *η*p^2^ = 0.002]).

In summary, we found that TMS over the left Ag selectively impaired RT performance for the retrieval of contextual information for highly coherent narratives (HC Cue Target conditions).

### 3.2. Reading Task

Reading times for the target passage showed a main effect of condition [F (2, 34) = 10.806, p < 0.001, *η*p^2^ = 0.389]. Post-hoc comparisons revealed that performance in the NC condition was overall slower than in the HC and LC conditions (p < 0.001 and p = 0.037, respectively). There was no significant RT difference for reading the target paragraph after the HC or LC contexts (p = 0.257). Finally, there was no significant effect of type of session [F (2, 34) = 0.551, p = 0.581, *η*p^2^ = 0.031], or significant a condition × session-type interaction [F (4, 68) = 0.231, p = 0.920, *η*p^2^ = 0.013] (see **Figure 4**).

**Figure 4.**
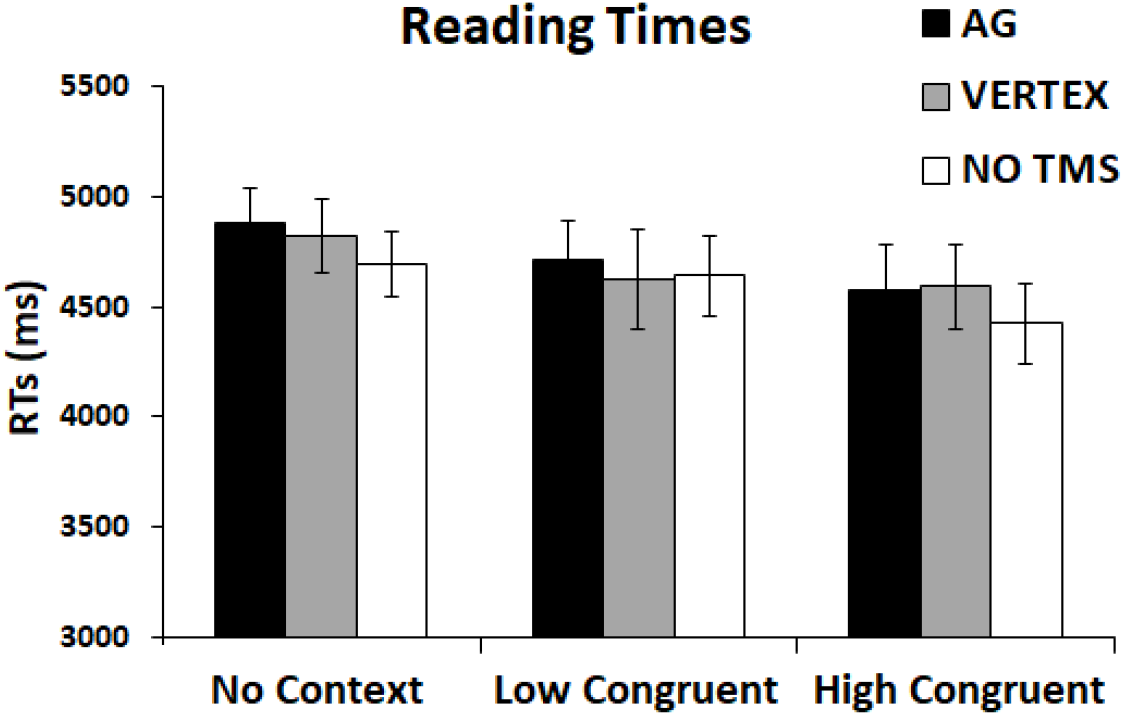
Reading task. Results for reading times for NC, LC and HC conditions. Error bars correspond to SEs.

## 4. Discussion

In the present study, we sought evidence for the hypothesis that the left Ag is critical for information buffering and context formation (Humphreys and Lambon Ralph, 2015, 2017; Branzi et al., 2019). This was tested by asking participants to read short narratives consisting of two sequential paragraphs (context and target) and by delivering TMS pulses over the left Ag between the context and target paragraphs, to disrupt online buffering and memory encoding of the narrative content (context and target integrated representation).

In the memory task, we measured RTs and proportion of correct responses to test the hypothesis that TMS-induced disruption of Ag activity during reading (encoding) would have had an effect on formation and therefore recall of integrated memory representation (context-target). We hypothesised that this effect would have been observed for HC conditions specifically, that is, when incoming information (target) matches the current knowledge-based schema (Speer and Zacks, 2005; Swallow et al., 2009; Humphreys and Lambon Ralph, 2015; van der Linden et al., 2017). In line with this hypothesis, we found that TMS delivered over left Ag made participants slower in retrieving context-related information during the memory task, only for coherent narratives (HC Cue Target conditions).

The results presented in this study fit well with the proposal that the core function of the left Ag would be that of an online automatic buffer (Humphreys and Lambon Ralph, 2015, 2017; Humphreys et al., 2019). This portion of the Ag, which is part of the default mode network (DMN), is positively engaged in semantic tasks, but only when these require context-dependent integration (Lerner et al., 2011; Simony et al., 2016; Baldassano et al., 2018; Branzi et al., 2019). Indeed, the very same parietal region is generally deactivated during semantic tasks (Visser and Ralph, 2011; Humphreys et al., 2015), casting some doubts about its key role in semantic integration. On the other hand, however, the left Ag is positively activated by tasks requiring episodic processing (Ramanan et al., 2018; Rugg and King, 2018).

One common aspect between the above mentioned semantic and episodic tasks is that they both likely require manipulation, retrieval and integration of episodic details (contextual who, what, when, where, why knowledge). In light of this observation, it is possible to relate to the involvement of the left Ag in our and previous studies to at least two mnemonic mechanisms. One would see the left Ag directly involved in the formation and representations of episodes (e.g., Shimamura, 2011; Bonnici et al., 2016, see the Contextual Integration Model proposed by Ramanan et al., 2018). This hypothesis fits with a series of findings that have shown that the left Ag’s activity is prominently observed at the end of an event, i.e., when all the information has been already presented (Humphries et al., 2007; Baldassano et al., 2017). Alternatively, the left Ag may be buffering the combination of past information (episodically retrieved) with newly presented information to update the growing contextual meaning. Not only this hypothesis accords with previous neuroimaging evidence (e.g., (Wagner et al., 2005; Vilberg and Rugg, 2008; Humphreys and Lambon Ralph, 2015, 2017; van der Linden et al., 2017; Branzi et al., 2019; Humphreys et al., 2019), but also with a various neuropsychological findings (for a review see Ramanan et al., 2018). In fact, patients with left Ag lesions are not amnesic like patients with hippocampal lesions on standard source and associative memory paradigms. Rather, damage to the left Ag seems to mainly diminish the confidence in recollection and the efficient online control of information (Simons et al., 2008; Simons et al., 2010). Accordingly, in the present study we show that TMS on the left Ag affects RTs, but not accuracy measures, therefore suggesting that disrupting the normal functioning of the left Ag does not cause the key information to be lost. Rather, TMS over the left Ag may have determined the formation of a less vivid and rich representation of a given episode, therefore making it less easy retrieve (see also Yazar et al., 2014).

Putting these results together, we propose that this DMN Ag region is more likely to be involved in the online buffer and use of episodically-recalled information rather than the coding and integration of this information per se (i.e., episodic and semantic representations might be coded in other regions, e.g., hippocampal regions and anterior temporal lobe, respectively). Still, the left Ag would be critical for the online buffering of internal and external information sources, which is a necessary precursor to the generation and the updating of context meaning.

It is worth to mention that, despite our expectation that TMS over left Ag will also impair behavioural performance during the reading task (and especially for HC conditions), our results did not support this prediction. In fact, we did not observe the expected disruption in reading times induced by Ag TMS. Although reading times have been previously taken as an index of the efficiency of online buffering processes during reading (e.g., Zwaan et al., 1995; Rinck and Weber, 2003; Speer and Zacks, 2005; Radvansky and Copeland, 2010), it is possible that the experimental design used here was not sensitive enough to detect the Ag TMS effect. In fact, in an attempt to allow enough time for our participants to memorise the content of the narratives, we may have allowed too much time for reading. This possibility is in accord with the fact that we did not replicate the LC>HC difference in reading times, previously observed in an fMRI study in which we allowed less time to respond (Branzi et al., 2019).

To conclude, our findings provide the first evidence of a causal role of the left Ag in information buffering and context formation during language processing and suggest that Ag activity observed in this and previous studies (e.g., Humphries et al., 2007; Baldassano et al., 2017; Branzi et al., 2019) is causally related to such processes. In fact, the TMS effect observed for HC Cue Target conditions only could not be explained as general episodic memory impairment.

Finally, in the memory task we also found slower RTs for LC Cue Target conditions as compared to HC Cue Target conditions. This result is in accord with previous studies that have shown that memory performance is affected by “event boundaries” (Swallow et al., 2009). In previous studies event boundaries were identified as changes in temporal, spatial and personal dimensions in the course of information processing. To our knowledge, this is the first study to demonstrate that also “semantic boundaries”, i.e., instances in which a change of semantic context occur, impair the formation and recall of integrated representations.

## Acknowledgments

this research was supported by a Postdoctoral Fellowship from the European Union’s Horizon 2020 research and innovation programme, under the Marie Sklodowska-Curie grant agreement No 658341 to FMB; an ERC Advanced Grant (GAP: 670428 - BRAIN2MIND_NEUROCOMP) and MRC Programme Grant (MR/R023883/1) to MALR and a Beacon Anne McLaren Research Fellowship (University of Nottingham) to JJ.

